# SARS-CoV-2 causes chronic lung inflammation and impaired respiratory capacity in aged Roborovski dwarf hamsters

**DOI:** 10.1101/2025.04.23.650177

**Authors:** Amirhossein Karimi, Carolin M Lieber, Kaori Sakamoto, Richard K Plemper

## Abstract

Roborovski dwarf hamsters are permissive for SARS-CoV-2 infection and progress to acute viral pneumonia with profound lung tissue injury, recapitulating hallmarks of severe COVID-19 in vulnerable patient groups such as older adults. In this study, we established dwarf hamster whole body plethysmography and assessed disease severity and propensity for long-term compromise of lung recovery from severe COVID-19-like disease in young, adult, and aged animals. Aged dwarf hamsters infected intranasally with variant of concern (VOC) omicron BA.4 experienced more severe clinical signs, carried a higher lung virus load, and had a greater risk of succumbing to infection. Resting airway hypersensitivity was transiently increased in aged, but not young, dwarf hamsters 3-4 days post infection (dpi). Pharmacologically induced respiratory distress revealed compromised lung capacity in animals of both age groups at peak disease. Aged animals showed impaired respiratory function for 45 days, mounted a weaker antiviral response, and developed chronic pneumonia with lasting tissue damage. Treatment of acute disease with approved antivirals, paxlovid-like nirmatrelvir+ritonavir or molnupiravir, prevented long-term respiratory sequelae in aged animals. Nirmatrelvir+ritonavir fully suppressed transient respiratory distress and mediated complete survival of aged animals. This study shows a high positive correlation between host age and SARS-CoV-2 disease severity in dwarf hamsters, establishes a model for chronic pneumonia with impaired respiratory capacity in at-risk hosts, and demonstrates benefit of antiviral therapy of acute disease for long-term respiratory health.

**Author Summary:** In the COVID-19 pandemic, the frequency of chronic respiratory insufficiency after acute SARS-CoV-2 infection was positively linked to patient age. Roborovski dwarf hamsters recapitulate hallmarks of life-threatening COVID-19 in at-risk patients that present with acute respiratory failure and prolonged respiratory incapacitation. In this study, we monitored disease progression and lung function in young and aged dwarf hamsters infected with a VOC omicron isolate and assessed the effect of antiviral treatment on long-term lung function. We established a strong correlation between host age and SARS-CoV-2 disease severity in dwarf hamsters, identified a high propensity of aged animals to develop chronic lung inflammation, and demonstrated a long-term loss of respiratory capacity in the subset of aged animals that survived the acute infection. Antiviral treatment suppressed the development of late sequelae and preserved lung function. These results have important implications for effective SARS-CoV-2 management in aged hosts at high risk of developing severe viral pneumonia with long-term impaired lung function.

## Introduction

Immunocompromised individuals and older adults are at greatest risk of developing develop severe COVID-19 after SARS-CoV-2 infection, which presents with acute viral pneumonia, major lung tissue injury, and high case-fatality rates [1]. Survivors frequently experience an extended recuperation phase with chronic pneumonia and prolonged, or permanent, reduction of respiratory capacity [2].

Several small-animal models have been developed to study SARS-CoV-2 pathogenesis, transmission, and late neuropathology sequelae. Mice are readily available and have low infrastructure demands but are not permissive for human SARS-CoV-2 strains without species adaptation, potentially confounding results and preventing direct analysis of relevant variants of concern (VOC) [3]. Ferrets have similar lung physiology to humans and support efficient SARS-CoV-2 shedding into, and transmission from, the upper respiratory tract without virus adaptation [3–5]. However, ferrets do not develop acute viral pneumonia after SARS-CoV-2 infection, thus poorly recapitulating severe disease in vulnerable human patient groups [3, 5]. Syrian golden hamsters have emerged as a model for acute infection of the upper and lower respiratory tract [3, 6, 7], and studies in aged versus young animals have revealed more severe lung lesions in older animals [8, 9]. However, the animals do not develop life-threatening SARS-CoV-2 pneumonia and mortality rates are low. The impact of Syrian golden hamster age on virus load remains furthermore controversial, since some studies reported a higher burden in older hamsters, whereas others found no such correlation [8–10]. Syrian golden hamsters have also been used to study respiratory complications after SARS-CoV-2 infection, but lung histopathology is transient and resolves by 14 days post infection (dpi) [7, 11]. Irrespective of animal age, longer-term studies in Syrian golden hamsters have remained limited to a 31-dpi window [7], which does not address whether chronic conditions may evolve.

None of the above models recapitulates hallmarks of life-threatening COVID-19 in at-risk patients that present with acute respiratory failure and prolonged respiratory incapacitation [3]. We have previously demonstrated that Roborovski dwarf hamsters are highly permissive for relevant SARS-CoV-2 VOC isolates without prior adaptation and develop acute viral pneumonia with profound lung tissue injury and high case fatality rates [12, 13]. Treatment of acute infection with approved antivirals alleviated clinical signs and altered disease outcome [12, 13]. This disease manifestation uniquely qualifies the model to assess the long-term benefit of antiviral therapy for the most at-risk patient populations under experimentally controlled conditions [12, 13].

During the pandemic, the frequency of chronic respiratory insufficiency after acute SARS-CoV-2 infection was positively correlated with patient age [14–16]. In this study, we assessed disease severity and propensity for long-term compromise of lung capacity after recovery from severe COVID-19-like disease in different age group dwarf hamsters representing young, adult, and aged animals, respectively. To non-invasively monitor the respiratory capacity of animals longitudinally, we developed conditions for dwarf hamster whole body plethysmography (WBP), which extracts respiratory parameters based on gas pressure changes due to respiratory activity of subjects in an airtight chamber (Fig 1a). The enhanced pause (Penh) is a dimensionless metric calculated from some of these parameters that is frequently interpreted as an indicator of respiratory distress [17–21]. Although the value of Penh as a measure for airway constriction is still a point of discussion [18, 22, 23], studies in SARS-CoV or influenza virus infected mice established a correlation between elevated Penh and the onset of clinical signs [20, 21], supporting the value of Penh as a quantitative measure of respiratory virus pathogenesis.

**Figure 1.**
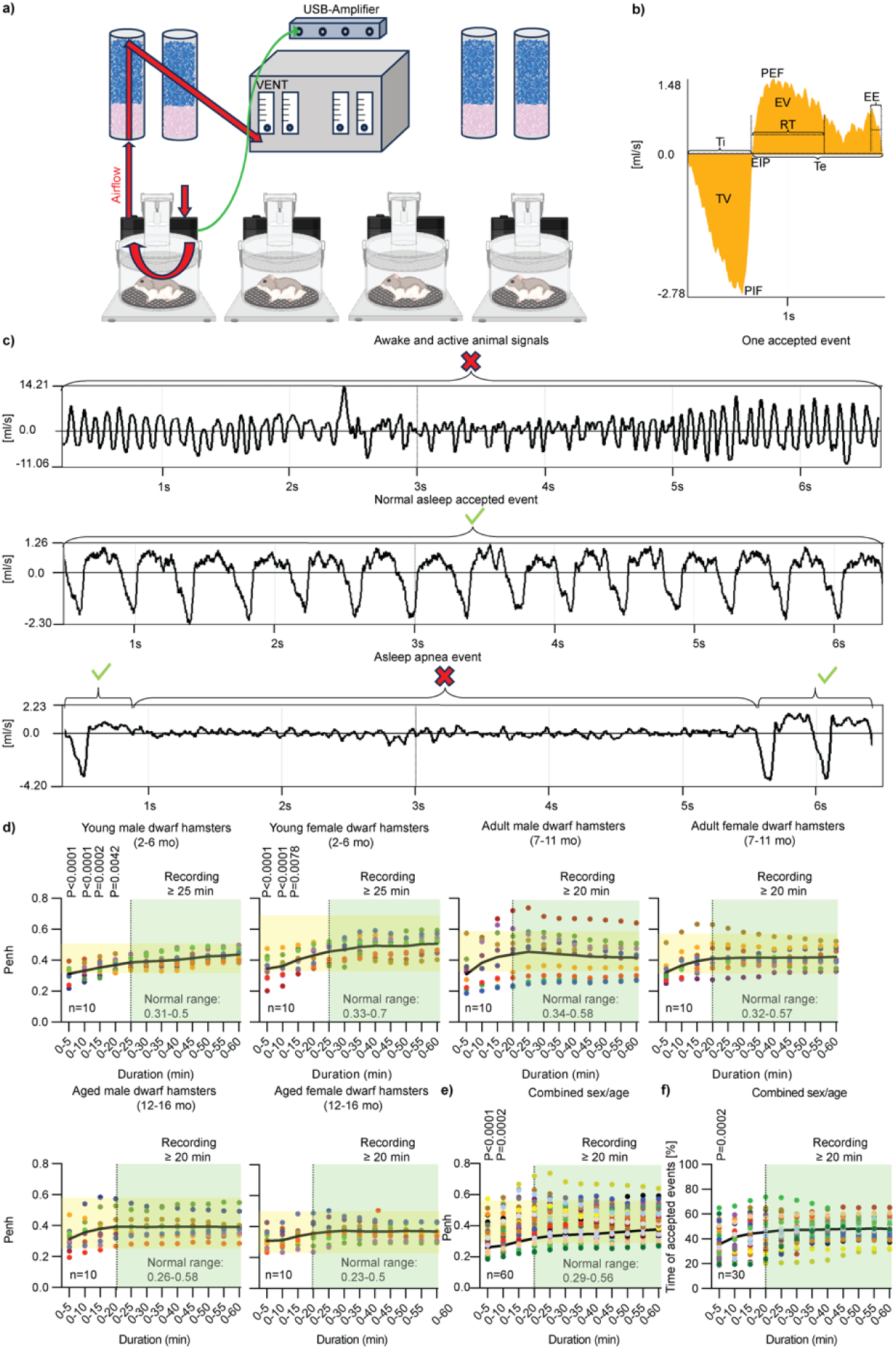
Whole-body plethysmography parameters in healthy Roborovski dwarf hamsters. **a)** Schematic of the WBP system, one central vent connected to four dry columns. **b)** Example of an accepted dwarf hamster breathing event. Inspiration (negative side of the y-axis) expiration (positive side). TV = Tidal volume (ml), Ti = Inspiratory time (ms), PIF = Peak Inspiratory Flow (ml/s), EIP = End Inspiratory Pause (ms), EV = expired volume (ml), Te = Expiratory Time (ms), RT = Relaxation Time (ms) the time required to exhale 30% of TV, EEP = End Expiratory Pause (ms). **c)** Active animal breathing (non-interpretable; top) and asleep breathing (interpretable; center), and a normal dwarf hamster apnea event (non-interpretable; bottom). **d)** Different time points of Penh value of WBP recording. Normal range (yellow highlight), plateau Penh mean (green highlight), n=10/sex and age group. Each color represents one animal. 1-way ANOVA with Dunnett’s multiple comparisons *post-hoc* test to determine analyze differences between individual time points and 60-minute recordings; P values <0.05 are shown. Penh normal ranges for each sex/age category are shown. **e)** Combined sex/age Penh value recording comparison at different time points (n=60). **f)** Accepted dwarf hamster breathing events at different time points compared to 60 minutes of WBP recording.

Our study revealed a strong positive correlation between host age and SARS-CoV-2 disease severity in dwarf hamsters, identified a high propensity of aged animals to develop chronic lung inflammation that lasted for over 45 dpi, and demonstrated a prolonged, and possibly permanent, loss of respiratory capacity in a subset of aged animals that survived the acute infection. Treatment of aged dwarf hamsters with paxlovid-like nirmatrelvir+ritonavir [12, 24, 25] or molnupiravir [13, 24, 25] alleviated clinical signs, completely suppressed the development of late sequelae, and preserved the full respiratory capacity of animals. These results have important implications for effective SARS-CoV-2 management in aged hosts at high risk of developing severe viral pneumonia.

## Results

We adapted a mouse whole body plethysmography (WBP) system to use with Roborovski dwarf hamsters, simultaneously monitoring respiratory function of four non-immobilized animals. During normal breathing, the respiratory response curve consists of distinct inspiration and expiration phases, each defined by the peak inspiratory and peak expiratory flow (PIF and PEF, respectively) of breath and the inspiration and expiration time (Ti and Te, respectively) (Fig. 1b). Penh is calculated from PEF, PIF, Te, and the time required for exhaling 70% of the breath volume (RT), according to the formula 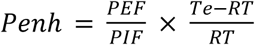 [18, 22].

### Baseline resting PenH range of different Roborovski dwarf hamster age groups

In contrast to mice, uninfected awake dwarf hamsters did not adhere to a regular breathing pattern, resulting in poorly defined and highly variable respiratory response curves that could not be interpreted for PenH calculation (Fig. 1c “awake and active”). Breathing curves became regular and interpretable, however, after animals fell asleep in the plethysmography chamber (Fig. 1c “asleep accepted”). An exception were periods of sleep apnea that occurred spontaneously in all animals, lasted for several seconds, but always resolved without intervention (Fig. 1c “asleep apnea”). Accordingly, most of the PenH calculations in this study are based on recordings that were taken of asleep animals at times of regular breathing.

To first establish PenH baselines for healthy dwarf hamsters, we sorted animals into six groups – young males or females (2-6 months of age), adult males or females (7-11 months of age), and aged males or females (12-16 months of age) – and subjected 10 animals of each group to 1-hour long WBP recordings (Fig. 1d). Normal resting PenH values did not differ statistically significantly between these groups, ranging from 0.23 (aged females) to 0.7 (young females). Young males showed the narrowest in-group spread, spanning from 0.31 to 0.5. Stable values were obtained in all groups when animals were recorded for at least 20 minutes (Fig. 1e), approximately 50% of which consisted, on average, of periods of acceptable events (Fig. 1f, S1 Fig, S1 Table).

### Effect of SARS-CoV-2 infection on resting PenH

SARS-CoV-2 infection of Roborovski dwarf hamsters causes acute lung injury, recapitulating disease presentation in vulnerable patient groups at high risk of severe COVID-19 [1, 2, 12–14]. To assess the effect of SARS-CoV-2 on respiratory fitness longitudinally, we intranasally infected adult dwarf hamsters in two subgroups per condition with 2,500 pfu/animal of variant of concern (VOC) BA.2, BA.4, or VOC delta. Peak lung virus load was determined in animals of one subgroup set at 3 dpi and the development of clinical signs, survival, and respiratory fitness was monitored in the other subgroups for a 14-day period (Fig. 2a). Animals in reference groups were mock-infected. Consistent with our previous work [13], VOC delta was highly pathogenic, causing major weight loss (Fig. 2b) and hypothermia (Fig. 2c), and most (3/4) animals succumbed within six days of infection (Fig. 2d). In contrast, clinical signs were mild or absent in animals infected with BA.2 or BA.4, and most animals recovered. Lung virus loads were similar in all 3 groups in the 10^5^-10^6^ pfu/g tissue range (Fig. 2e). Resting PenH was statistically significantly elevated in the 3-4 dpi window in dwarf hamsters infected with VOC delta, reaching peak values of 2-3 when animals approached predefined endpoints (Fig. 2f). However, resting PenH did not deviate from the normal range in animals of the VOC omicron BA.2 and BA.4 groups.

**Figure 2.**
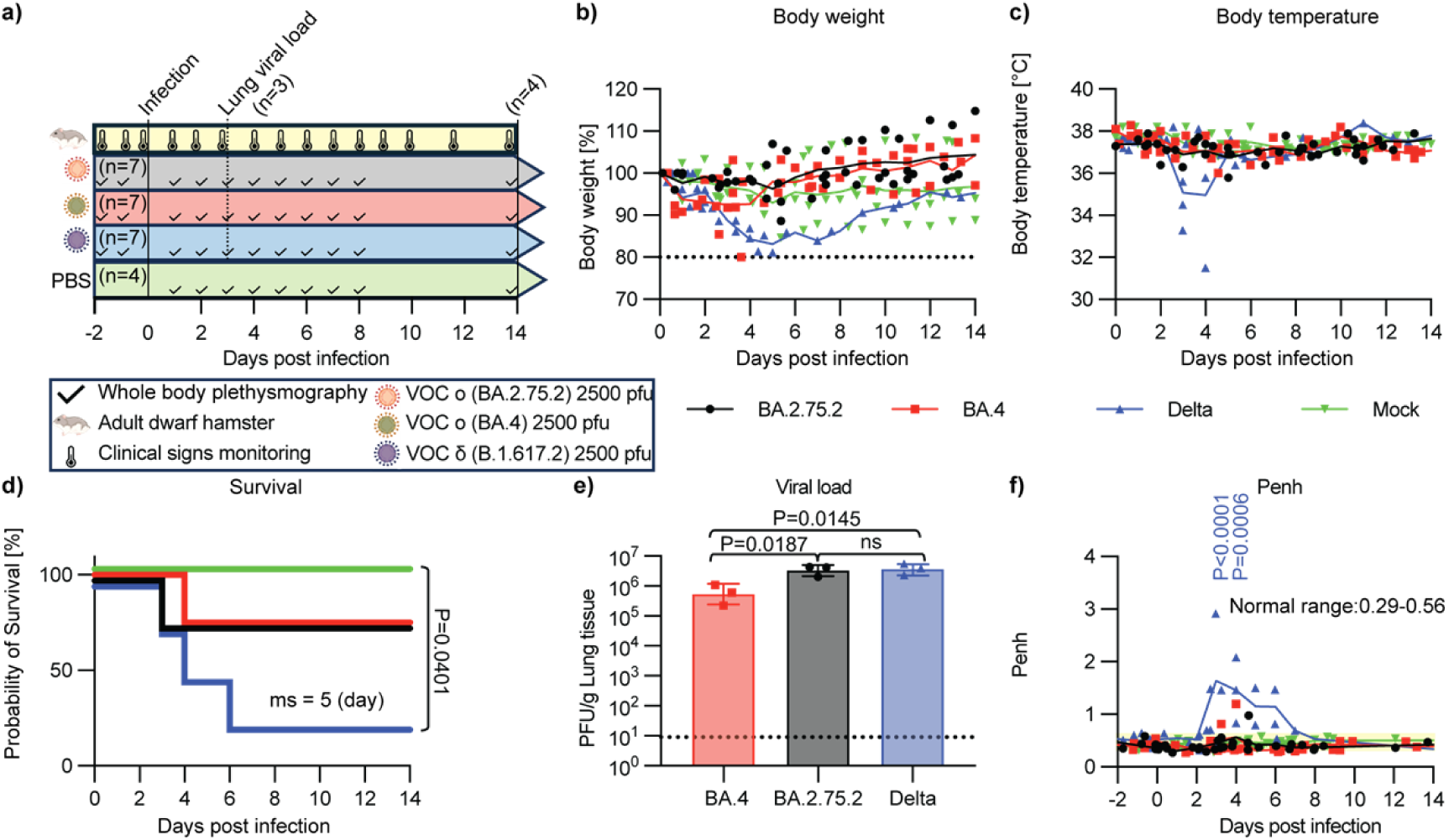
Significant increase in Penh in dwarf hamsters infected with SARS-CoV-2 VOC δ. **a)** Three groups of adult dwarf hamsters (n=7/group) infected with different SARS-CoV-2 variants of concern and mock-infected animals (n=4). Three dpi, lungs of infected hamsters (n=3) were harvested for viral load determination. All remaining animals were euthanized 14 dpi. **b,c)** Body weight (b) and temperature (c) of all groups were monitored for 14 dpi (weights were normalized to initial % weights and all data are shown staggered). **d)** Probability of survival. Three animals in VOC δ infected, and one each infected with VOC o (BA.2.75.2) and (BA.4) succumbed to infection (median survival shown). Log-rank (Mantel-Cox) test to determine the significance between different groups (only P<0.05 is shown). **e)** Lung viral load 3 dpi. Columns represent geometric data means ± geometric SD, symbols show individual animals. One-way ANOVA with Tukey’s multiple comparisons *post-hoc* test; P values <0.05 are shown. **f)** Calculated Penh post infection. Penh values among different groups were compared to mock-infected dwarf hamsters; 2-way ANOVA with Dunnett’s multiple comparisons *post-hoc* test; P values <0.05 are shown.

These results corroborated previous findings [13] of high pathogenicity of VOC delta in adult dwarf hamsters that presents with acute lung injury and indicated that solely late-stage severe viral pneumonia results in appreciable respiratory distress in the model.

### Stress PenH conditions in Roborovski dwarf hamsters

Inhaled methacholine causes constriction of the airways and is used clinically in pharmacological bronchial challenge tests [26, 27]. To establish conditions for Roborovski dwarf hamster stress-PenH, we exposed male and female hamsters of the three age groups to increasing concentrations of aerosolized methacholine ranging from 0.3-80 mg/ml for 30 seconds, followed by WBP measurement (Fig. 3a). Animals in all groups were exposed to aerosolized saline, followed by WBP reference recording before methacholine challenge. Tolerability limits were 80 mg/ml in young animals and adult females, and 40 mg/ml in adult males and aged animals (Fig 3b). In all groups, stress PenH values deviated first from baseline when methacholine concentrations exceeded 4 mg/ml (S2 Fig). Dwarf hamster resting respiratory function was fully restored 15 minutes after methacholine exposure (S2 Table). Accordingly, all subsequent stress PenH studies were carried out with 4 mg/ml methacholine, inhaled for 30 seconds, followed by a 15-minute recovery period.

**Figure. 3.**
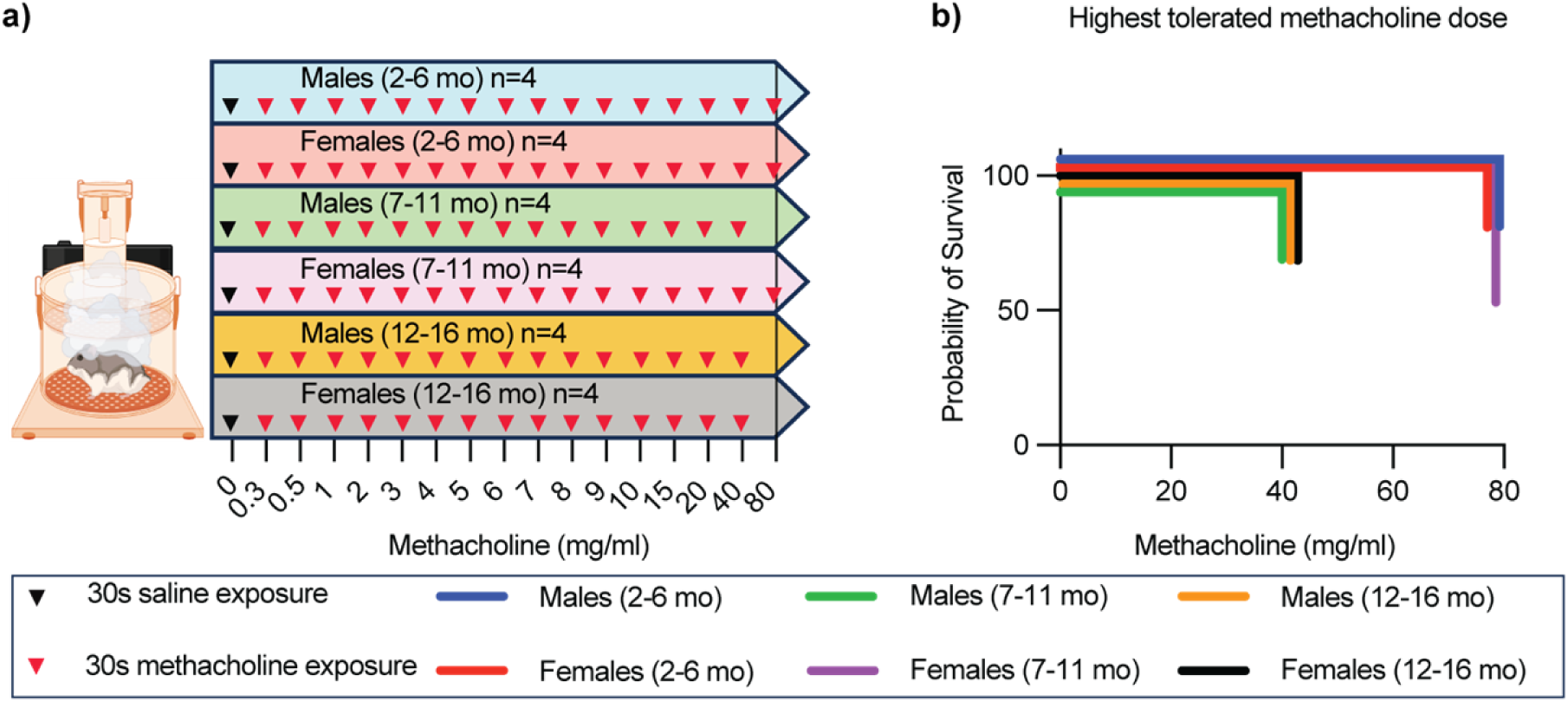
Highest tolerated methacholine dose in healthy Roborovski dwarf hamsters. **a)** Study schematic. Sixteen different doses of methacholine were nebulized in different age/sex dwarf hamsters (n=4/group). Maximum of 3 doses/day; 30 seconds of exposure of each dose followed by 15-minute WBP recording. **b)** Probability of survival after exposure. Adult (green) and aged (orange) males and aged females (black) dwarf hamsters are shown to be more susceptible to higher doses of methacholine exposure than younger animals (blue and red) and adult females (purple).

To assess a possible effect of isoflurane anesthesia and intranasal infection on resting and stress PenH readout, we next subjected dwarf hamsters to WBP after anesthesia alone, or anesthesia followed by inoculation with 50 μl (25 μl per nare) sterile saline (S3a Fig). Reference animals were not anesthetized. None of these manipulations triggered changes in resting PenH, but stress PenH was transiently increased beyond the normal range for 2 (anesthesia only) to 3 (anesthesia plus PBS inoculation) days (S3b,c Fig).

### Aged dwarf hamsters are at greater risk of developing severe COVID-19-like disease

Older adult COVID-19 patients were highly vulnerable to developing viral pneumonia with acute lung tissue injury that required hospitalization [2, 14]. To examine the effect of age on disease severity in the dwarf hamster model, we infected young and aged animal groups with 1,000 pfu of VOC omicron BA.4, which was confirmed as a non-lethal inoculum amount in the young group (Fig. 2d). Clinical signs were monitored daily as before and resting and stress PenH determined daily for the first 14 dpi and in 2-3-day intervals thereafter (Fig. 4a). All young animals survived the infection without developing clinical signs, whereas 50% (6/12) of aged dwarf hamsters succumbed after experiencing profound weight loss and hypothermia (Fig. 4b-d). Mean lung virus load determined 3 dpi in a subset of animals from either age group was statistically significantly higher in aged animals, exceeding that detected in young dwarf hamsters by approximately one order of magnitude (Fig. 4e).

**Figure 4.**
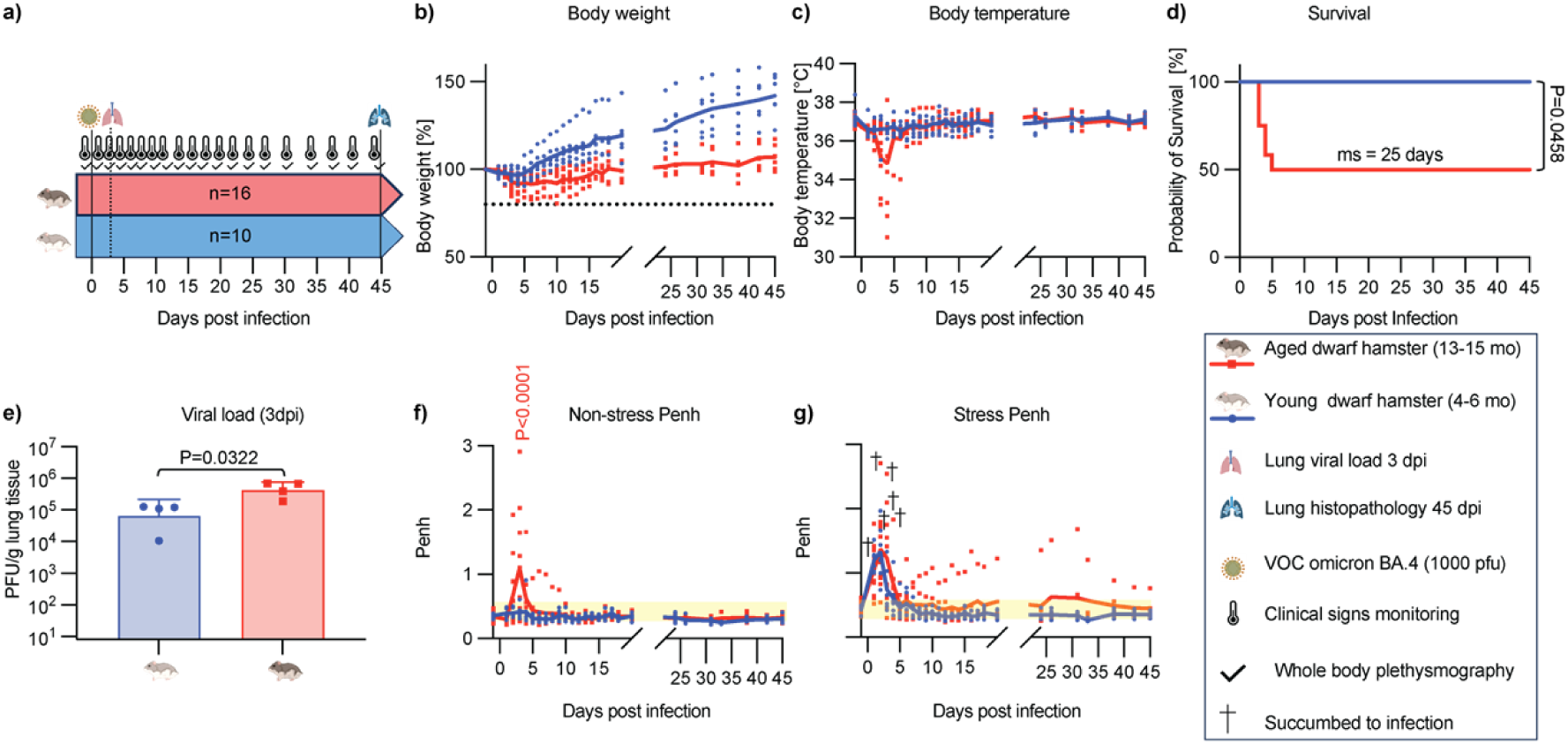
Aged Roborovski dwarf hamsters are highly susceptible to SARS-CoV-2 infection with long-lasting respiratory complications. **a)** Study schematic. Animals were infected intranasally with 10^3^ pfu of VOC omicron BA.4. Lungs were harvested for viral load determination at 3 dpi (dotted line) (n=4/group). Clinical signs (black thermometer icon) were monitored for 45 dpi and WBP (black check mark) was measured. Lungs of the remaining animals were harvested for histopathology assessment (blue lung symbol) at 45 dpi study end. **b,c)** Body weight (b) and temperature (c) in aged (red) and younger (blue) dwarf hamsters at 45 dpi. **d)** Probability of survival shown with calculated median survival. Log-rank (Mantel-Cox) test to determine the significance between two groups; P value is shown. **e)** Lung viral load in young and aged dwarf hamsters at 3 dpi. Columns represent geometric means ± geometric SD, symbols show individual animals. Unpaired t-test to assess significance between the groups. P value is shown. **f,g)** Non stress-Penh (f) (30s normal saline, 0.9% sodium chloride, exposure and 15-minute WBP), and stress-Penh (g) evaluations; 30 seconds methacholine (4 mg/ml) exposure and 15-minute WBP. Combined sex/age Penh normal range, 0.29-0.56 (yellow highlighted) and last Penh reading before death (black cross) are marked; 2-way ANOVA with Bonferroni’s multiple comparisons *post-hoc* test to determine significant changes in Penh between groups; P value <0.05 is shown.

### Prolonged elevated stress PenH after recovery of aged animals from SARS-CoV-2

Resting PenH of aged dwarf hamsters peaked in the acute disease phase and most animals with PenH values exceeding 1.0 succumbed to the infection (Fig. 4f). By 10 dpi, resting PenH of all surviving aged animals returned to baseline and resting PenH of young animals remained unremarkable throughout the study. In contrast, stress PenH showed statistically equivalent initial peaks in both age groups, followed by full recovery in young animals by 7 dpi (Fig. 4g). However, stress PenH remained elevated in several surviving aged dwarf hamsters until study end 45 dpi, suggesting prolonged compromise of respiratory function.

Histopathology of lungs extracted at study end from the 6 surviving aged dwarf hamsters and the young animals revealed signs of inflammation in 4 of 6 aged animals (Fig. 5a,b). Three aged animals presented with fulminant interstitial pneumonia and foamy macrophages were present in alveoli of another aged dwarf hamster. In contrast, lung tissue appearance in all young animals was unremarkable, resembling that of uninfected animals. No difference in tissue appearance between young and aged animals was detected in the uninfected reference group.

**Figure 5.**
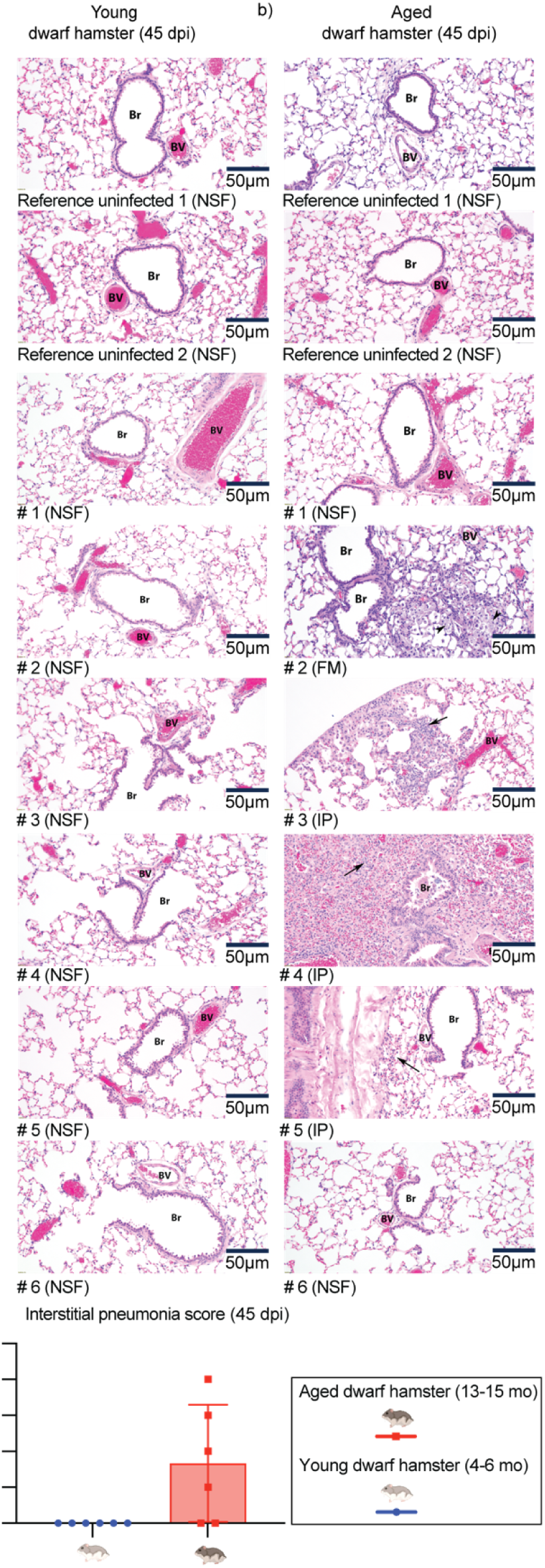
Alveolar histopathology showed long-term sequelae in aged animals. **a)** H&E staining of dwarf hamster lung tissue; magnification 20×. Each # represents a different animal. NSF: no significant findings, FM: foamy macrophages (black arrowhead); IP: interstitial pneumonia (black arrows). **b)** Interstitial pneumonia scores of survivors 45 dpi. Columns represent data means ± SD, symbols show individual animals.

These results demonstrate that aged dwarf hamsters are at significantly greater risk of developing severe, life-threatening COVID-19-like disease than young animals. The majority of aged survivors experienced prolonged pulmonary inflammation coinciding with compromised respiratory function.

### Antiviral treatment preserves respiratory capacity in aged dwarf hamsters

To assess the potential therapeutic benefit of antiviral therapy in aged hosts, we administered two therapeutics approved for human use, paxlovid (nirmatrelvir+ritonavir) and molnupiravir, to assess the effect of treatment on changes in respiratory capacity during, and after recovery from, SARS-CoV-2 infection. Aged dwarf hamsters were inoculated intranasally with 1,000 pfu of VOC omicron BA.4, followed by oral treatment with nirmatrelvir (250 mg/kg)+ritonavir (41.5 mg/kg) or molnupiravir (250 mg/kg) first initiated 12 hours after infection and continued b.i.d. for 7 days (Fig. 6a). Animals were monitored daily as before, lung tissue extracted from a subset of 4-5 animals per condition 3 dpi, and WBP carried out daily for the first 2 weeks, then in 3–4-day intervals until 45 dpi. Vehicle-treated animals transiently lost weight (Fig. 6b) and developed hypothermia (Fig. 6c), and 50% of animals reached predefined endpoints within 7 dpi (Fig. 6d). Clinical signs were greatly alleviated or absent in animals of both treatment groups. However, only treatment with nirmatrelvir+ritonavir resulted in complete survival, likely reflecting the poor pharmacokinetic performance of molnupiravir in dwarf hamsters [13]. Lung virus load was statistically significantly lower in animals of the nirmatrelvir+ritonavir group than in either vehicle or molnupiravir-treated animals (Fig. 6e).

**Figure 6.**
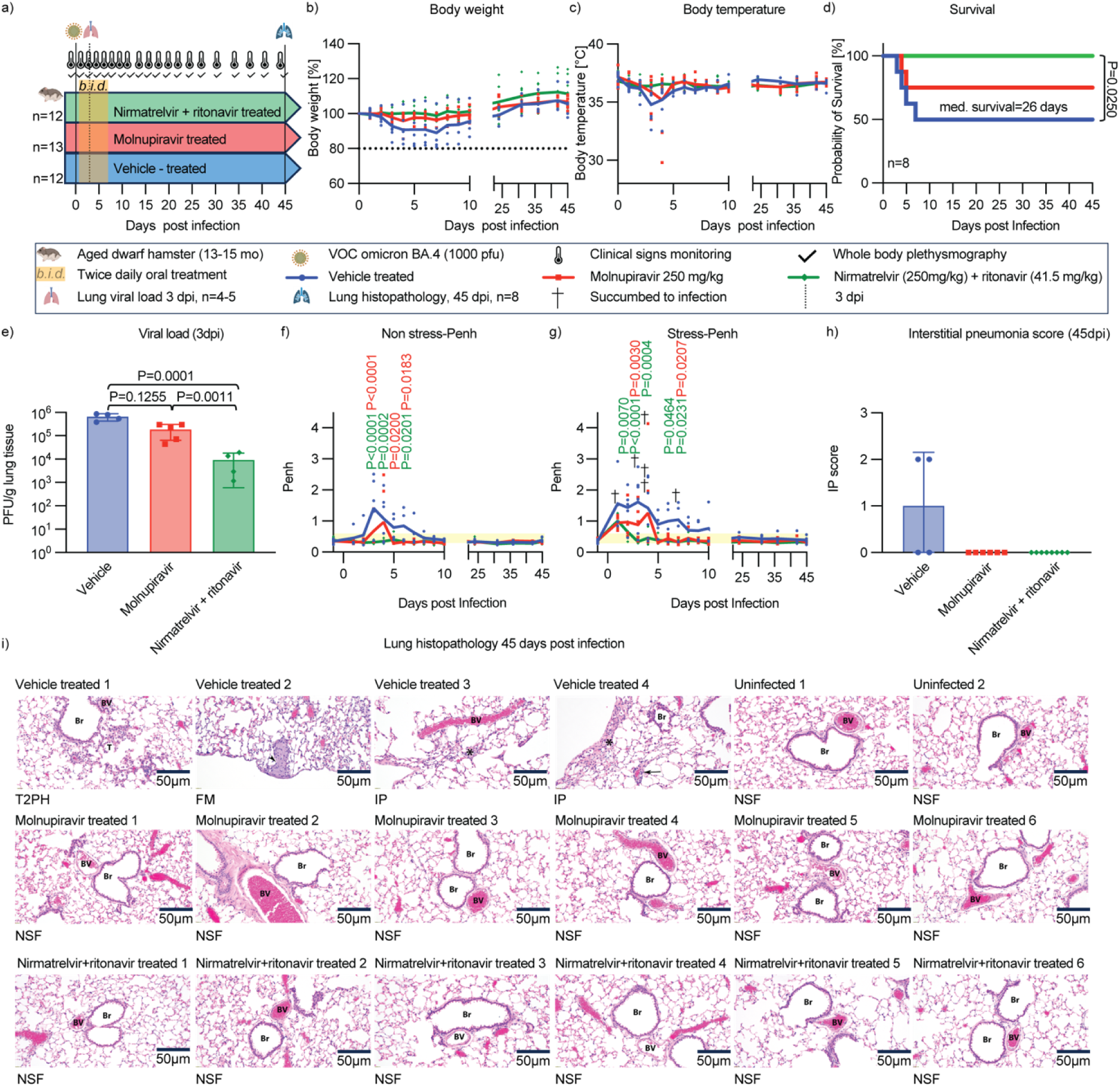
Antiviral treatment preserved lung function in aged dwarf hamsters infected with SARS-CoV-2. **a)** Study schematic. Aged dwarf hamsters were infected intranasally (n=13/group) with BA.4; 10^3^ pfu/animal. Treatment with molnupiravir (250 mg/kg) (red), nirmatrelvir (250 mg/kg)+ritonavir (41.5 mg/kg) (green), or vehicle (blue) (b.i.d. regimen) was initiated 112 hrs after infection. Clinical signs were monitored throughout the study. **b,c)** Body weight (b, normalized to initial weight %) and temperature (c). Symbols represent individual animals, lines connect data means. **d)** Probability of survival. Log-rank (Mantel-Cox) test to calculate median survival. **e)** Lung virus load 3 dpi. Columns represent geometric means ± geometric SD, symbols show individual animals. 1-way ANOVA with Tukey’s multiple comparisons *post-hoc* test; P values <0.05 are shown. **f,g)** Non stress (f, saline exposure) and stress-Penh (g, methacholine 4 mg/ml exposure) comparisons between different groups. Penh normal range (0.29-0.56) (yellow highlights). Final stress Penh for individual animals prior to death (indicated by black crosses). 2-way ANOVA with Dunnett’s multiple comparisons *post-hoc* test; P values <0.05 are shown. **h)** Interstitial pneumonia score of survivors at study end. Columns represent means ± SD, symbols show individual animals. **i)** H&E staining of lung sections of animals that survived until study end (45 dpi) and of uninfected references. No significant findings (NSF), T2Ph: type 2 pneumocyte hyperplasia (black T), FM: foamy macrophages (black arrowhead); IP: interstitial pneumonia (black arrow and asterisk); magnification 20×. Each picture shows the lung of a different animal.

Resting PenH was unremarkable in nirmatrelvir+ritonavir-treated dwarf hamsters (Fig. 6f), resembling the profile of untreated young animals. Molnupiravir reduced respiratory distress compared to vehicle-treated dwarf hamsters, but individual animals in the treatment group that became moribund experienced greatly elevated resting PenH at 4 dpi. Stress PenH showed an initial peak 1 dpi in all groups (Fig. 6g). However, peak height was lower, and the duration of elevated stress PenH shorter, in animals of both treatment arms compared to the vehicle group, and none of the treated animals experienced prolonged respiratory distress (Fig. 6g,h). Lung histopathology of all treated animals was unremarkable at study end (45 dpi), whereas all animals in the vehicle group showed lesions consistent with inflammatory injury (Fig. 6i). These results demonstrate that early-onset treatment of SARS-CoV-2 infection of aged dwarf hamsters suppressed prolonged lung inflammation and prevented tissue damage, providing major long-term therapeutic benefit through preserving respiratory health.

### Inflammatory responses in aged versus young dwarf hamsters

We determined mRNA expression levels of a selected panel of cytokine genes in lung tissues of young and aged animals daily 1-4 dpi with VO C omicron BA.4, relative to those in uninfected animals of the respective age group to probe for differences in the quality of the antiviral responses (Fig. 7a). A final subgroup of only aged hamsters was sampled at 45 dpi to explore possible effects of chronic infections. Expression profiles revealed three distinct trends: cytokines with consistently higher relative expression in young than in old animals; cytokines with approximately equal response pattern; and cytokines with greater upregulation in aged than in young dwarf hamsters (Fig. 7d). Notably, major pro-inflammatory cytokines such as TNF-α, IL-12, and IL-17 were found in the first group, indicating induction of a robust response in young, but not aged, animals (Fig. 7b). We did not notice statistically significant differences in induction of IL-6, IFN-β, or IFN-γ responses between the groups (Fig. 7c). However, anti-inflammatory IL-10 was statistically significantly upregulated in aged dwarf hamsters compared to young animals. At 45 dpi, all measured expression rates had returned to homeostasis in aged animals. These results are consistent with higher initial virus load and prolonged infection of aged dwarf hamsters.

**Figure 7.**
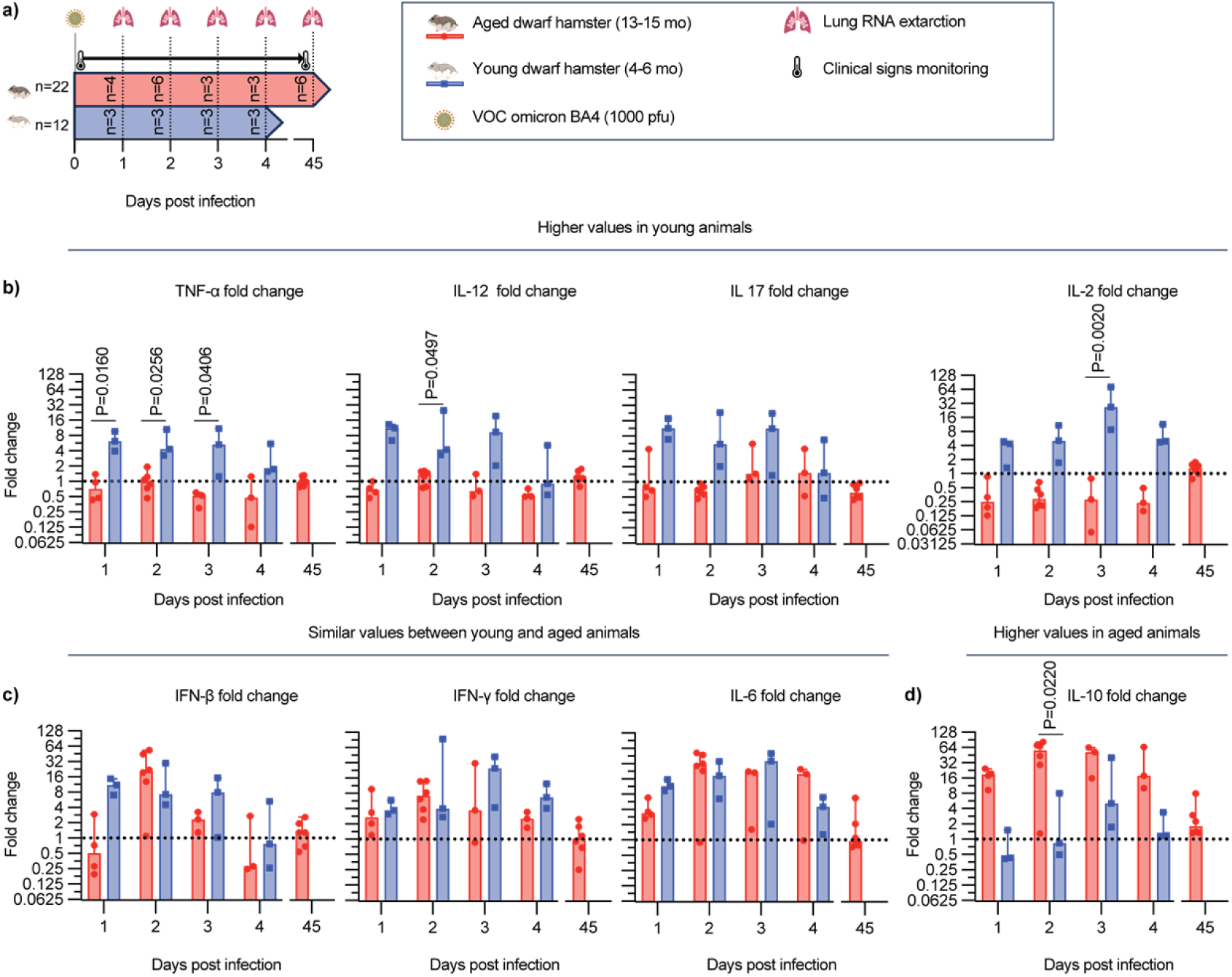
Relative expression profiles of selected cytokines in young versus aged dwarf hamsters. **a)** Study schematic. Only aged hamsters were sampled at 45 dpi. **b-d)** qRT-PCR to quantify changes in lung mRNA levels relative to those of uninfected animals. Results were sorted by expression profile into higher relative induction in young animals (b), parity (c), and higher induction in aged animals (d). Symbols show results for independent repeats (individual animals) determined in two technical repeats each, columns represent group means. 2-way ANOVA with Bonferroni’s multiple comparison *post-hoc* test; P values <0.05 are shown.

## Discussion

In this study, we applied WBP to Roborovski dwarf hamsters for non-invasive monitoring of respiratory distress after SARS-CoV-2 infection, which required establishing baseline breathing parameters for the species. Due to their high activity level and metabolic rates [28], assay set-up and baseline parameters available for other rodent species, such as mice [18, 19] or Syrian golden hamsters [29], were not transferable to dwarf hamsters. Non-anesthetized animals could not be restrained without major distress, which would have confounded measurements, and WBP plots of awake dwarf hamsters were devoid of interpretable breathing patterns. Because the animals are nocturnal, however, WBP of asleep hamsters emerged as a viable solution. Highly reproducible respiratory capacity parameters were obtained for healthy dwarf hamsters in three relevant age groups, young, adult, and aged, each subdivided into male and female animals. Variations in key WBP parameters between groups were minor, resulting overall in a robust reference dataset for this species that is available for query in future work.

Infection of different dwarf hamster age groups with SARS-CoV-2 and subsequent longitudinal monitoring of animals through the acute disease, recuperation, and post-infection stages supported three major conclusions:

*i) Dwarf hamster age is a major correlate for pathogenicity and overall disease outcome.* We detected a statistically significant correlation between animal age and severity of COVID-19-like disease in Roborovski dwarf hamsters, including the magnitude of clinical signs, peak lung virus loads at acute infection, and prospect of survival. Analogous studies in Syrian golden hamsters likewise revealed a trend towards increased clinical signs in older animals [8, 9]. However, outcomes of studies exploring the effect of age on lung virus load in the Syrian golden hamster model are mixed, since some reports describe increased virus burden in the upper and lower respiratory tract [8, 30], whereas others concluded that animal age had no effect on virus replication [9]. Consistent across studies, lung pathology was increased in aged Syrian golden hamsters due to impaired tissue repair [7–9, 30], a slower and weaker innate antiviral response [8], and an impaired adaptive response characterized by elevated levels of regulatory T cells and neutrophils and reduced quality of neutralizing antibodies [8, 9].

Neutrophils in particular have been implicated in both viral clearance early after infection and subsequent contribution to tissue damage [7–9]. Although a paucity of validated reagents prevented direct analysis of specific dwarf hamster immune cell populations, increased SARS-CoV-2 pathogenicity and profound lung tissue injury strongly support that diminished innate and adaptive immunity similarly underlies enhanced disease in aged dwarf hamsters. In contrast to the Syrian golden hamster model, however, case-fatality rates are high in dwarf hamsters infected with SARS-CoV-2 [13, 31]. Our WBP results demonstrate that death was due to respiratory failure, recapitulating the presentation of at-risk human patients with severe SARS-CoV-2 pneumonia [1, 14, 32].

*ii) Surviving aged animals are at significantly greater risk of experiencing chronic lung lesions and prolonged compromise of respiratory capacity*. Whereas Syrian golden hamsters fully recover from SARS-CoV-2 infection within a 2-week window without appreciable lasting adverse effects [7, 9], late sequelae including prolonged periods of reduced respiratory capacity occur frequently in humans [2, 32]. Following infected animals longitudinally over 7 weeks, we observed that the majority of aged dwarf hamsters surviving the acute disease stage developed chronic respiratory lesions. Lung histopathology showed residual interstitial pneumonia, aggregates of foamy macrophages, or type 2 pneumocyte hyperplasia in 4 of 6 animals, which coincided with impaired respiratory capacity. These results illuminate a propensity of dwarf hamsters to develop post-COVID-19-like sequelae that phenocopy hallmarks of slow recuperation after clearance of acute SARS-CoV-2 infection described in human patients.

*iii) Treatment of acute infection substantially improves the prospect of full recovery of aged animals*. We have demonstrated previously that dwarf hamsters present with viral pneumonia within less than 24 hours of infection with SARS-CoV-2 [13]. Thus, at 1 dpi animals are at the transition stage to complicated disease, which human patients suffering from severe COVID-19 typically reach in the second week after infection [14, 33, 34]. Based on these distinct kinetics of disease progression, we believe initiation of antiviral treatment 12 hours after infection of dwarf hamsters represents a realistic mimic of therapeutic intervention started at the transition point to severe disease. Testing the two orally bioavailable antivirals that first received approval for treatment of COVID-19, paxlovid-like nirmatrelvir+ritonavir and molnupiravir, our study demonstrates that treatment of acute SARS-CoV-2 infection alleviates clinical signs and eliminates chronic lung inflammation in aged hosts, preserving respiratory capacity. However, only nirmatrelvir+ritonavir statistically significantly reduced lung virus load and mediated complete survival of aged dwarf hamsters. This result was unexpected, since we had observed comparable efficacy of both drugs in previous studies with ferrets, mice, and adult dwarf hamsters [12, 13, 35]. Considering the equivalent long-term therapeutic benefit of preserving respiratory capacity of both treatments, we believe this difference reflects that the pharmacokinetic properties of molnupiravir in aged dwarf hamsters are less favorable than in younger animals or non-rodents [4, 12, 13]. Most likely, tissue exposure of the triphosphate form of the bioactive molnupiravir metabolite is capped in aged dwarf hamsters at a level below that required to sufficiently suppress virus replication to achieve complete survival. Indeed, a link between host age and altered nucleotide metabolism due to deregulated nutrient sensing is well documented [36]. Because of the limited number of aged dwarf hamsters available, however, we could not further pursue this question experimentally.

The maximal parallel processing capacity of the WBP system used in this study was four chambers, which dictated maximal animal group sizes per condition examined. However, substantially larger number of animals were measured to establish the normal reference values for each age group. Other study limitations derived from the outbred nature of Roborovski dwarf hamsters, which enhances animal-to-animal variability of results. Since dwarf hamsters used in this study were not bred under controlled conditions, variation between individual animals is exacerbated by unknown disease history of individual animals prior to sourcing and an uncertainty factor in documented animal age of ± 1 month. We have increased group sizes in several studies and/or repeated entire study arms with separately sourced and aged dwarf hamsters to ensure reproducibility of our conclusions, but the issue is inherent to this animal model and can only be partially mitigated. However, outbred animals with distinct disease history that were raised under open range rather than controlled husbandry conditions and were of similar, albeit not identical, age are far more representative of human patient groups than inbred research animals [1, 2, 14, 16, 37, 38]. While experimental variability is increased, confidence in the physiological relevance of emerging, statistically significant trends identified in this study is, therefore, likewise heightened.

In conclusion, this study mapped baseline WBP breathing parameters for Roborovski dwarf hamsters of different age groups, demonstrated increased susceptibility of aged dwarf hamsters to develop acute, life-threatening COVID-19-like viral pneumonia after SARS-CoV-2 infection, and revealed that the majority of aged animals recovering from acute infection developed chronic lesions with prolonged, and possibly permanent, reduction of respiratory capacity. This model recapitulates hallmarks of severe COVID-19 in older adults and provides a robust system to assess the benefits of antiviral therapy in at-risk patients under experimentally controlled conditions. Full preservation of respiratory capacity in this proof-of-concept study with approved antivirals confirmed that treatment of acute disease fully mitigates late respiratory sequelae frequently associated with SARS-CoV-2 infection of aged hosts.

## Materials and Methods

### Study design

Young, adult, and aged male and female dwarf hamsters were used in *in vivo* experiments. Animals of different ages and sexes were infected intranasally with distinct SARS-CoV-2 VOC, and pathogenesis of the disease was followed short and long-term using WBP. Treatment efficacy of authorized antivirals against SARS-CoV-2 (molnupiravir and nirmatrelvir+ritonavir) was evaluated based on the WBP parameters after infection. Treatments started 12 hours post-infection and continued twice daily for 7 dpi in survival groups and 2.5 dpi in viral load assessment groups. Oral gavage of molnupiravir was performed in 1% methylcellulose, nirmatrelvir in 0.5% methylcellulose + 2% Tween80, and ritonavir in 22% ethanol. Vehicle-treated groups received 1% methylcellulose. Viral loads were determined by plaque assay from whole lung tissues 3 dpi. Clinical signs were monitored twice daily for 7 dpi and once daily for 7-14 dpi.

### Cells and viruses

African green monkey kidney cells VeroE6-TMPRSS2 (BPS Bioscience #78081) were cultured at 37°C with 5% CO_2_ in Dulbecco’s Modified Eagle’s Medium (DMEM) with 7.5% heat-inactivated fetal bovine serum (FBS). These cells were used to perform plaque assay for lung viral load evaluation. SARS-CoV-2 VOC delta (B.1.617.2) and omicron (lineages BA.2.75.2 and BA.4) were maintained under biosafety level 3 (BSL-3) conditions. Virus stock titers were confirmed via plaque assay prior to *in vivo* usage.

### Plaque assay

Ten-fold serial dilutions were generated from the initial sample in DMEM medium which contained antibiotics (Gibco). Serial dilutions were transferred to 12-well plates seeded with VeroE6-TMPRSS2 cells 24 hours earlier at a density of 2.5 × 10^5^ cells/ml (0.5 ml/well). Infected plates were incubated for 1 hour at 37 °C, 5% CO_2_. After the incubation period, the inoculum was replaced with 2.4% Avicel (FMC BioPolymer), which was mixed with 2×DMEM (1:1 ratio) and plates were incubated for 3 days at 37°C with 5% CO2. After incubation, Avicel was removed and cells were washed with PBS, fixed, and stained with 10% neutral formalin buffer and 1% crystal violet in 20% ethanol.

### Roborovski dwarf hamsters

Roborovski dwarf hamsters were purchased from Dierengroothandel Ron Van Der Vliet, Netherlands at an estimated age of 8-10 weeks. Upon arrival, animals were housed in a biosafety level 1 (ABSL-1) facility at Georgia State University for, minimally, a 14-day acclimation period prior to study start.

### Whole-body plethysmography

The WBP system was purchased from SCIREQ scientific respiratory equipment. The unit consists of four chambers, each of which can accommodate one dwarf hamster for simultaneous recordings. Bias flow was set to 0.5 lpm for all experiments. Air exiting the chambers passed through drierite desiccant columns (W.A.Hammond Drierite Co) prior to venting. Average temperature and humidity in the system were set to 70 F and 50% RH. For all WBP chambers, flow threshold was set to 0.75 ml/s, Ti range to 60-1000 ms, Te range to 80-1000 ms, TV range to 0.04-10 and F range to 10-600 bpm. Measurements for RT were obtained at 30% of TV and for EIP/EEP at 5% of PIF/EV. The beginning of inspiration was set at 20% of PIF. Breathing rate was computed from Ti + Te. A maximum of 40% deviation from inspiration/expiration volume triggered rejection of the measurement.

### WBP recording parameters

Normal 1-hour WBP recordings from 10 animals (young, adult, and aged/male and female) were randomly selected to assess average Penh in different time frames compared to PenH resulting from 60-minute recordings (n=10/group and n=40/universally combined group). For each animal, average Penh was calculated 5, 10, 20, 25, 30, 35, 40, 45, 50, 55, and 60 minutes from the beginning of the recording. Twenty animals of different sex and age were selected to determine the minimum required time to calculate Penh for each subgroup. WBP times of accepted events was calculated for each animal 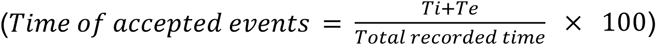 within the same time frames.

### Normal ranges of different WBP parameters for Roborovski dwarf hamsters

A total of 235 dwarf hamsters (young, adult and aged, male and female) were used to determine the WBP normal ranges for different respiratory parameters. One-hour/day WBP recordings were performed for each animal. The first WBP recording was considered acclimation of the animal to the WBP chamber, and the data received from the 2^nd^ day or the average of recordings of days 2 and 3 were utilized for normal range determination and categorized based on the age/sex of the animals in addition to the universally established normal ranges.

### Intranasal infection of dwarf hamsters

Dwarf hamsters were anesthetized with ketamine 50-75 mg/kg and dexmedetomidine 0.5 mg/kg, followed by inoculation intranasally with 500-5000 pfu in 50 μl sterile PBS (25 μl per nare). Anesthesia was reversed using Antisedan at 1 mg/kg body weight.

### Tissue sample collection

Animals were euthanized through prolonged 5% isoflurane inhalant followed by cervical dislocation of anesthetized animals. Whole lungs were removed, weighed and homogenized using a tube containing glass beads and 300 μl sterile PBS with 2× antibiotics-antimycotics cocktail (Gibco). Samples were bead blasted in 3 cycles (30 seconds each) with 30-second interruptions at 4°C. Subsequently, samples were centrifuged for 10 minutes at 3,000 rpm and 4°C. Following centrifugation, supernatants were aliquoted and stored at −80°C until virus titration.

### Lung cytokine profiling

Approximately 30 mg of lung tissue (extracted from the same lobe for each animal) was homogenized and total RNA extracted using the RNeasy mini kit (Qiagen). cDNA corresponding to message was synthesized from total RNA extracts using oligo-dT primers and Invitrogen SuperScript III reverse transcriptase. Fast SYBR Green Master Mix (Applied Biosystems) was used to perform real-time PCR with validated primers against selected dwarf hamster cytokines or glyceraldehyde-3-phosphate dehydrogenase (S3 Table). Threshold cycle (ΔΔC_T_) values were calculated relative to uninfected reference animals to determine the expression fold-changes.

### Aerosolized methacholine exposure

A designated nebulizer (SCIREQ scientific respiratory equipment) was connected to each WBP chamber for methacholine delivery. A total of 26 healthy dwarf hamsters in 6 groups (different ages/sexes as previously explained) were randomly selected to determine the tolerability of methacholine exposure in this species. All animals were first exposed to saline (sterile 0.9% sodium chloride solution) for 30 seconds, followed by 15-minute WBP recording. The same time frame of exposure and WBP was then applied for methacholine doses of 0.3, 0.5,1, 2, 3, 4, 5, 6, 7, 8, 9, 10, 15, 20, 40, and 80 mg/ml. All exposures were induced at 15% of the delivery cycle. A maximum of 3 doses of methacholine/day were tested in dwarf hamsters after 15 minutes of daily baseline recording.

### Histopathology

The lungs, after being infused with 10% neutral-buffered formalin, were submerged in that solution. After 24h, the formalin was replaced with 100% ethanol for 48h. Paraffin was used to embed the fixed lung samples, and they were subsequently sectioned, processed, and stained with hematoxylin and eosin (H&E). Histopathology scoring and lung tissue assessment were performed by a board-certified veterinary pathologist who was blinded to the study groups. Interstitial pneumonia was scored based on the thickness of alveolar septa due to infiltration by leukocytes (1 = 1 leukocyte thickness, 2 = 2 leukocytes thick, 3 = 3 leukocytes thick, 4 = 4 leukocytes thick) in the most severely affected area.

### Statistical analyses

Unpaired t-tests were applied for comparison between two independent groups. Sample sets containing more than two group were analyzed using 1-way or 2-way ANOVA depending on the number of variables, followed by Tukey’s, Dunnett’s, or Bonferroni’s multiple comparison *post-hoc* tests as specified in figure legends. Statistical significance was defined as p-values ≤ 0.05.

### Software

Statistical analyses were performed using the Prism 10.4.1 (GraphPad) software package. Graphs were assembled in Adobe Illustrator 2024, study schematics were generated in Biorender. The iox 2.10.8.38 package was used to analyze WBP raw recording data.

### Ethics statement

The Guide for the Care and Use of Laboratory Animals of the National Institutes of Health was applied to animal work, in full compliance with the Animal Welfare Act Code of Federal Regulations. *In vivo* studies in dwarf hamsters were approved by the Georgia State University Institutional Animal Care and Use Committee (IACUC) under protocols A21019 and A24009. Work with SARS-CoV-2 strains and the experimental analysis of infectious materials and animal tissues was approved by the Georgia State University Institutional Biosafety Committee (IBC). All cell culture and *in vivo* studies were performed in BSL-3/ABSL-3 containment facilities at Georgia State University.

## Acknowledgements

We thank M.G.Natchus and G.R.Painter for providing EIDD-2801 (molnupiravir) used in this study. This work was supported, in part, by public health service grant U19 AI171403 from the NIH/NIAID (to RKP and GRP). All experimental work in this study supported by this award was completed at or before March 24, 2025.

## Author Contributions

A.K., data curation, formal analysis, investigation, methodology, writing – original draft; C.L., data curation, formal analysis, investigation, writing – review & editing; K.S., investigation, writing – review & editing; R.K.P., conceptualization, formal analysis, funding acquisition, project administration, supervision, validation, writing – original draft

## Competing Interests

The authors declare that no competing interests exist.

## Supporting Information Captions

**S1 Table.** Normal ranges of WBP-parameters in Roborovski dwarf hamster groups representing different ages and sexes.

**S2 Table.** Minimum time required for a full recovery after 30-second methacholine exposure.

**S3 Table.** Primer sequences of Roborovski dwarf hamster cytokines used for qPCR.

**S1 Figure. Percentage of accepted events in different time frames of WBP recordings. a-f)** Uninfected dwarf hamsters of different ages and sexes (n=5/group). Each color shows an individual replicate, black lines connect sample means. Significant differences were evaluated by 1-way ANOVA with Dunnett’s multiple comparison *post-hoc* test, comparing the means of each time point to the same group means after 60-minute recording; P values <0.05 are shown.

**S2 Figure. Methacholine dose-range finding for stress Penh measurements. a-f)** Penh (mean) comparison after exposure to different dose-levels of methacholine. Statistical analysis with 1-way ANOVA with Dunnett’s multiple comparison *post-hoc* test; P values <0.05 are shown. Penh normal ranges for each age/sex group are highlighted in yellow. Biological repeats (independent animals) are shown in different colors (n=4).

**S3 Figure. Anesthesia and mock infection transiently increase stress Penh. a)** Study schematic. Mock infection intranasally with 50 µl PBS. Animals in the anesthesia-only group received ketamine + dexmedetomidine (IP), animals in the reference group did not receive any treatment (n=4/group). **b,c)** Saline solution or 4 mg/ml methacholine were nebulized to determine non-stress (b) or stress-Penh (c), respectively. Yellow bars denote normal ranges; 2-way ANOVA with Dunnett’s multiple comparison *post-hoc* test; P values <0.05 are shown.

## Notes

### Competing Interest Statement

The authors have declared no competing interest.

